# Comprehensive catalog of dendritically localized mRNA isoforms from sub-cellular sequencing of single mouse neurons

**DOI:** 10.1101/278648

**Authors:** Sarah A. Middleton, James Eberwine, Junhyong Kim

## Abstract

RNA localization to neuronal dendrites is critical step for long-lasting synaptic potentiation, but there is little consensus regarding which RNAs are localized and the role of alternative isoforms in localization. Using independent RNA-sequencing from soma and dendrites of the same neuron, we deeply profiled the sub-cellular transcriptomes to assess the extent and variability of dendritic RNA localization in individual hippocampal neurons, including an assessment of differential localization of alternative 3’UTR isoforms. We identified 2,225 dendritic RNAs, including 298 cases of 3’UTR isoform-specific localization. We extensively analyzed the localized RNAs for potential localization motifs, finding that B1 and B2 SINE elements are up to 5.7 times more abundant in localized RNA 3’UTRs than non-localized, and also functionally characterized the localized RNAs using protein structure analysis. Finally, we integrate our list of localized RNAs with the literature to provide a comprehensive list of known dendritically localized RNAs as a resource.

## Introduction

Neurons require local protein synthesis within the dendrites to produce long-lasting synaptic potentiation (Aakalu et al. 2001; Eberwine et al. 2001; Job and Eberwine 2001). In order for this local synthesis to occur, mRNAs must first be transported to the dendrites. Although RNA localization and local translation have been studied for over 20 years, including initial sanger sequencing of isolated single dendrite RNA (Miyashiro, Dichter, and Eberwine 1994; Crino and Eberwine 1996), a more detailed and thorough analysis is required to generate a consensus set of dendritically localized RNAs. Surprisingly, the advent of high-throughput sequencing has not greatly improved matters: of three recent RNA-seq studies of dendritically localized RNA (Cajigas et al. 2012; Ainsley et al. 2014; Taliaferro et al. 2016), only 1% of the identified RNAs overlapped between all three studies (44 of 4,441). Although these differences can be partly attributed to differences in sample origin, organism, and experimental protocol between each study, these examples nonetheless point to a need for further studies to understand the full range and variability of dendritic RNAs.

There are several major challenges in profiling the dendritic transcriptome: (1) cleanly separating the somatic and dendritic compartments so that they can be profiled separately, (2) differentiating transcript variation (e.g., alternative 3’UTRs) in addition to localization, and (3) accounting for single cell variation in both somatic expression and dendritic localization. Given that substantial gene expression heterogeneity has already been observed on the whole-neuron level (Dueck et al. 2015), it would not be surprising if there is variability of localization across cells, as was found in an early single dendrite sanger sequencing study (Miyashiro, Dichter, and Eberwine 1994). In addition, localization variability in neurons may arise from the use of alternative 3’UTR isoforms. Neurons uniquely express a large number of extended 3’UTR isoforms that are conserved between human and mouse (Miura et al. 2013), and one possibility is that a subset of these 3’UTRs contain dendritic localization signals. A few specific examples of differentially localized 3’UTR isoforms have already been characterized (Miura et al. 2014), such as BDNF (An et al. 2008; Liao et al. 2012). Taliaferro *et al.* recently surveyed this phenomenon on a larger scale in brain-derived cell lines and cortical neurons and identified hundreds of cases of differential localization of alternative 3’UTR isoforms (Taliaferro et al. 2016).

Here, we expand upon these earlier studies by performing simultaneous RNA-sequencing of the somatic and dendritic compartments of single neurons from primary cultures to allow for a direct contrast of the dendritic transcriptome with its parent soma and to enable the assessment of heterogeneity of localization across neurons. Using this single neuron sub-cellular sequencing approach, we identify dendritically enriched RNAs on both the gene and isoform levels, including several of the recently identified neuron-enriched distal 3’UTR extensions (Miura et al. 2013). We identify a total of 2,225 candidate dendritic RNAs, including 298 that showed differential localization of 3’UTR isoforms that was consistent across the individual cells. Using structure-and sequence-based computational techniques, we extensively annotate these dendritic RNAs to explore their functions and identify possible motifs involved in dendritic targeting.These new computational models provide a library of testable predictions that will help dissect the molecular mechanism of dendritic localization and dendritic RNA function. Finally, we integrate our list of dendritic genes with the current literature, producing a definitive list of dendritic RNAs that have been observed to date in high-throughput studies.

## Results

### Identification of dendritically localized RNAs

To compare the RNAs present in dendrites and somas of individual neurons, we manually separated the dendrites and soma of primary mouse hippocampal neurons using a micropipette (Miyashiro, Dichter, and Eberwine 1994) and performed RNA-sequencing on each subcellular fraction such that we obtained the subcellular transcriptomes of the same cell (Fig. 1A). We note that the axon is generally small at this culture stage (~5% the volume of the dendrites) with a thin gauge (< 1uM) and has a flush axon hillock which is easily distinguishable from a dendrites graded hillock. Thus, we do not expect the axon to be harvested in our procedure, and any axon that was collected would not make up a large fraction of the isolated dendrite samples. A total of 16 individual neurons were collected (32 soma and dendrite samples). Extracted RNA was amplified using the aRNA procedure (Morris, Singh, and Eberwine 2011; Van Gelder et al.1990; Eberwine et al. 1992) and sequenced to an average depth of 25 million reads per sample. Somas generally contained a wider variety of transcripts than their corresponding dendrites, with an average of 9,206 and 5,827 genes identified in each compartment respectively. As expected, the genes represented in the dendrites were largely a subset of the soma-expressed genes of the same cell (Fig. 1B). All soma and dendrite samples expressed housekeeping genes and neuronal marker genes at high levels, especially pyramidal cell markers such as *Grin1, Mtap2*, and *Neurod6*, with little expression of other brain cell type markers (Fig. 1C).

**Figure 1.**
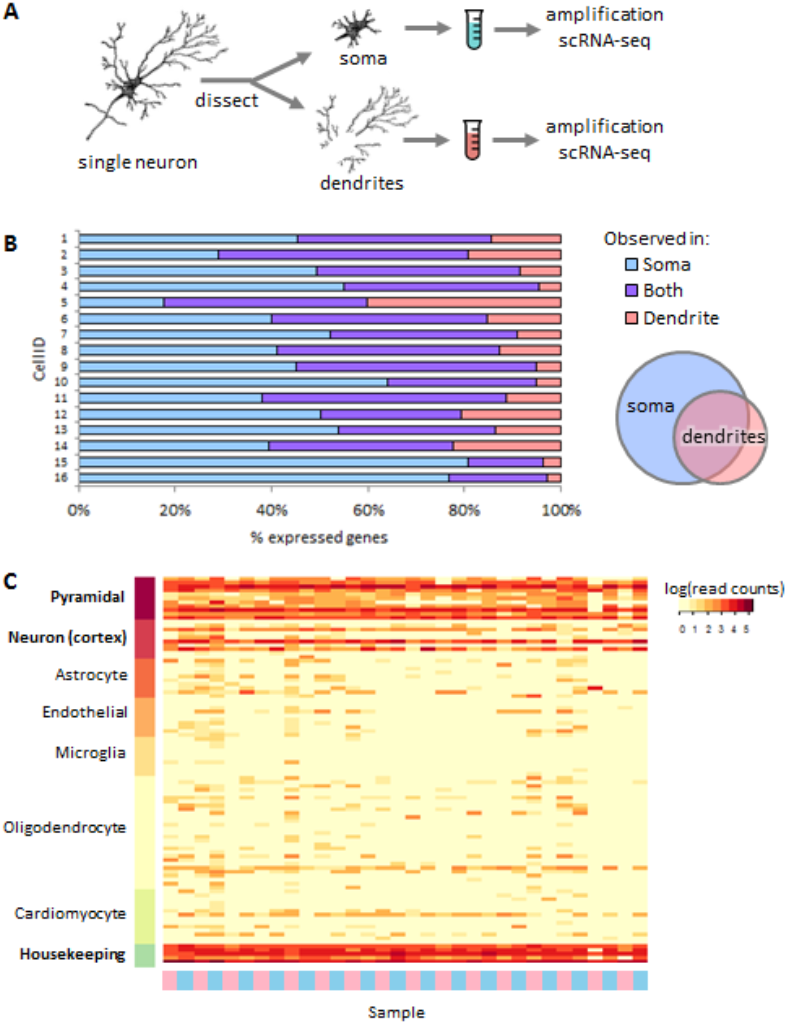
Sub-single cell profiling of soma and dendrite RNA. (A) Isolated single neurons were dissected to separate the soma and neurites, which were collected into separate tubes for amplification and RNA-sequencing. (B) Overlap of expressed genes (≥10 reads) between soma and dendrites from the same original cell. Each horizontal bar shows the results from a single neuron. The Venn diagram depicts the general relationship between the somatic and somatic transcriptome of the same cell. (C) Marker gene expression for several brain cell types. Samples (columns) are indicated as either dendritic samples (pink) or soma samples (blue). Cardiomyocte markers are included as a control cell type that is electrically active but unrelated to brain cells.

To identify potentially localized RNAs, we used DESeq2 (Love, Huber, and Anders 2014) to perform a differential expression analysis using a paired design, where soma and dendrites of the same original cell were directly compared. DESeq2 reported 3,811 genes significantly more highly expressed in somas and 387 genes significantly higher in dendrites (FDR corrected p ≤ 0.05) (Fig. 2A). Given their relatively higher expression in dendrites compared to soma, these 387 genes are likely to be actively localized, and we therefore refer to them as localized RNAs. The localized RNAs were strongly enriched for GO terms related to translation and mitochondria, consistent with previous reports (Ainsley et al. 2014; Francis et al. 2014; Taliaferro et al. 2016), whereas the somatic RNAs were enriched for functions related to the nucleus, including RNA splicing and chromatin organization (Fig. 2B and Supplemental Table S1). Notably, there was no significant enrichment among these localized genes for terms specifically related to plasticity or synaptic function.

**Figure 2.**
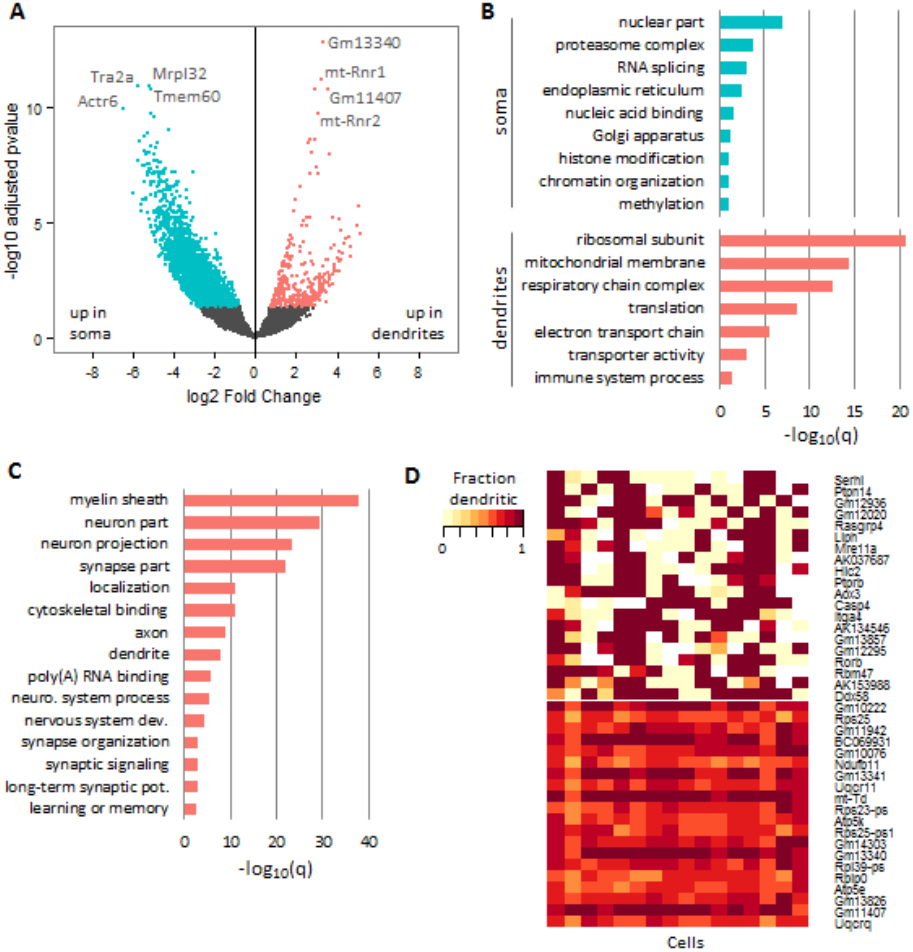
Differentially expressed genes between soma and dendrites. (A) Differentially expressed genes in soma (blue) and dendrites (pink). (B) Selected GO terms enriched in the soma and dendrites (deDend) based on the differential expression analysis. (C) Selected GO terms enriched in the consDend genes. (D) Heatmap showing the dendritic read fraction for the top 40 genes (rows) with the highest and lowest variability of localization. Each column represents a single cell.

Differential expression analysis may not identify all localized RNAs because not all localized RNAs are expected to have higher expression concentration in the dendrites than the soma. This may be particularly relevant when expression is profiled at the single cell level, since factors such as bursting transcription and variable rates of localization can lead to high variability in the relative amounts of RNA in each compartment at the time of collection. Therefore, we additionally identified RNAs that were consistently present in the dendrites across the profiled cells, since these RNAs are likely to have important dendrite function even if they are not differentially at higher concentration in the dendrites compared to the soma. We found 1,863 RNAs in at least 90% of the dendrite samples, which included well-characterized localized RNAs such as *Actb, Bdnf, Calm1, Dlg4, Grin1*, and *Map2*. To differentiate from the 387 differentially expressed genes described above, we refer to this set as the constitutive dendritic (consDend) RNAs, and the previous set as the differentially expressed dendritic (deDend) RNAs. The consDend RNAs covered many of the same ontology functions as the deDend RNAs, such as mitochondria and translation, but additionally were strongly enriched for a large number of synaptic and localization-related GO terms (Fig. 2C and Supplemental Table S1). The consDend RNAs also contained a large number of genes with the GO annotation “myelin sheath”, which is unexpected given that this term is normally associated with axons. However, closer examination showed that this term includes genes with a wide variety of other functions (Supplemental Table S1), and the consDend list does not contain myelin basic protein (*Mbp*). Overall, the differences between the deDend and consDend lists suggest that at the single cell level, RNAs with important dendritic and synaptic functions are often not localized to the point of having higher expression concentration in the dendrites relative to the soma, but are nonetheless consistently present in the dendrites at a lower level.

Single cell analysis also allows us to examine the variability of localization across cells. For each of the 387 deDend RNAs, we calculated the variation of localization across cells based on the variance of the dendritic read fraction (defined as the number of dendritic reads divided by the sum of the dendritic and somatic reads for each cell). The top 40 genes with the highest and lowest localization variability are shown in Figure 2D. The high variability genes had lower median total-cell expression (dendritic + somatic reads) than the low variability genes (76.6 and 415.7 reads, respectively), and it should be noted that differences in expression level can potentially contribute to observed variability in single cell experiments. From a biological perspective, low variability of localization suggests a gene is localized by a constitutive mechanism and is needed in constant supply in the dendrites, whereas high variability suggests more dynamic localization mechanisms which may be activated in response to stimuli. The genes with the highest variability of localization included several enzymes (*Serhl, Ptpn14, Liph, Mre11, Aox3, Casp4, Ddx58*), most of which do not currently have a defined dendritic function, although mutations in *Mre11* have been previously associated with Ataxia-telangiectasia-like disorder 1 (Stewart et al. 1999). These high variability genes also showed more “all-or-nothing” localization than the low variability genes, with most cells having a dendritic read fraction of close to either zero or one (Fig. 2D). Genes with the least variable localization included components of the ubiquinol-cytochrome c reductase complex (*Uqcrq, Uqcr11*), ATP synthase complex (*Atp5e, Atp5k*), and ribosomal subunits (*Rplp0, Rps25*), some of which in humans have been implicated in schizophrenia and schizoaffective disorder (Arion et al. 2015). These results give further support to the idea that genes involved in respiration and translation are needed in constant supply in the dendrites, and suggest that this might be accomplished by a constitutive localization mechanism that is relatively constant across cells.

### Differential localization of 3’UTR isoforms

Given the potential importance of alternative 3’UTR usage in dendritic localization, we sought to better define genes that have 3’-isoform-specific dendritic localization in primary neurons. As a result of the aRNA single cell RNA amplification process (Morris, Singh, and Eberwine 2011; Van Gelder et al. 1990; Eberwine et al. 1992), the majority of our sequencing reads map within 500nt of a 3’ end (Fig. 3A), and we thus have high coverage of these regions for identifying expressed 3’UTR isoforms. We quantified the expression of individual 3’ isoforms based on the last 500nt of each isoform, merging any 3’ ends that were closer than 500nt into a single feature. Individual cells widely expressed multiple 3’ isoforms per gene, with somas showing slightly more alternative expression than dendrites on average (1.26 and 1.13 expressed 3’UTR isoforms per gene, respectively; Fig. 3B). When multiple isoforms were expressed, one isoform tended to be dominant, making up ~85% of the gene reads on average in both compartments. To compare differential isoform representation between soma and dendrite, we limited the considered 3’UTR isoforms to only the top two most highly expressed isoforms per gene, which accounted for the vast majority of reads in most genes. The top two isoforms were labeled “proximal” (the more 5’ isoform) or “distal” (the more 3’ isoform), and isoform preference for each gene in each sample was summarized as the fraction of reads mapping to the distal isoform (distal reads divided by distal plus proximal reads), which we refer to as the distal fraction (DF). We focused our analysis only on multi-3’UTR genes that had at least 10 total reads in both the soma and dendrites of at least five cells, which resulted in 3,638 considered genes. We note that alternative 3’UTRs can be generated by two distinct mechanisms: alternative splicing, which generates alternative last exons (ALEs), or alternative cleavage and polyadenylation, which generates tandem UTRs (Fig. 3C). Therefore, we split our set of multi-3’UTR genes into ALE and tandem groups based on the relationship between the designated proximal and distal 3’UTR for that gene. ALEs made up the majority of the considered multi-3’UTR genes (3,108 ALE versus 530 tandem).

**Figure 3.**
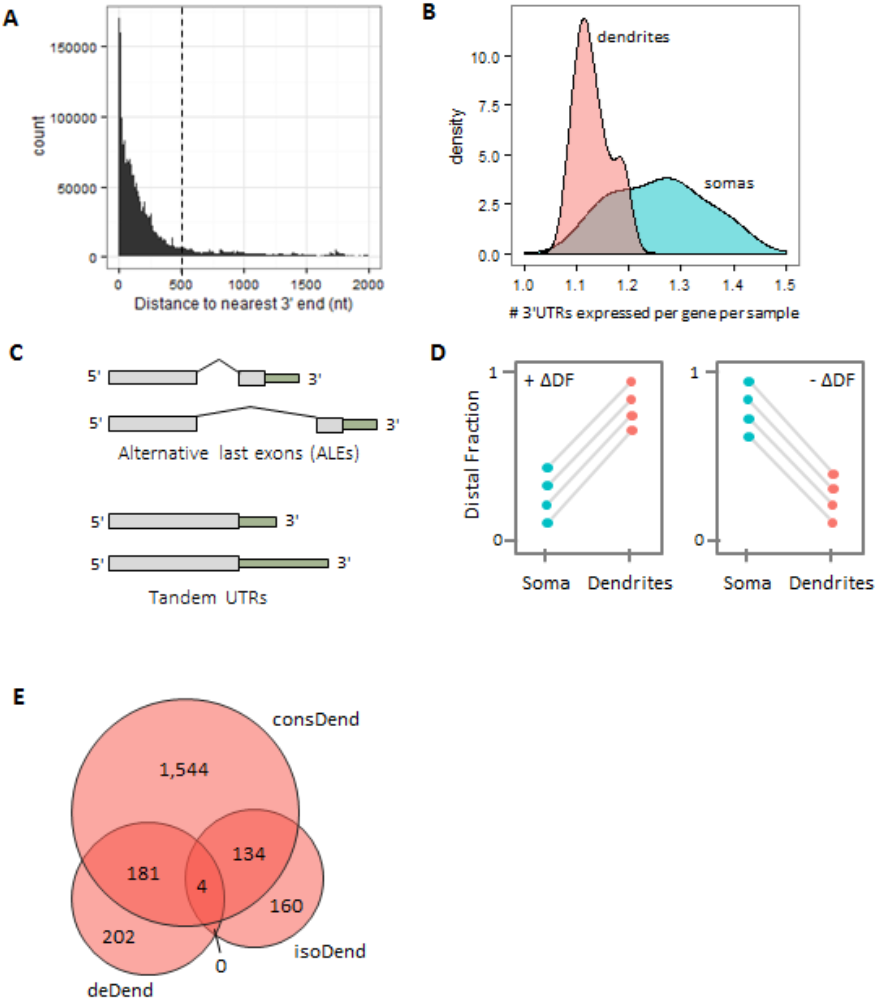
Alternative 3’UTR isoform usage in neurons. (A) Distribution of distance from read ends to the nearest gene 3’ end. Most reads are within 500nt of the nearest end (dotted line). (B) Distribution of the number of 3’UTRs expressed per gene per sample in dendrite samples (pink) and soma samples (blue). (C) Definition of ALEs and Tandem UTRs. (D) Theoretical examples of genes with consistent changes in distal fraction (ΔDF) across cells, shown as paired plots. Somas and dendrites from the same original cell are shown connected by a line. Consistently positive (left) or negative (right) ΔDF indicates differentially localized isoforms between the two compartments. (E) Overlap between the three sets of dendrite-localized genes (gene-level, resident, and isoform-level).

To identify 3’UTR isoforms that are differentially localized to dendrites, we looked for genes that had consistent patterns of isoform preference across our cells. That is, we looked for cases where the change in distal fraction (ΔDF; defined as DF_dendrite_–DF_soma_ and calculated separately for each soma-dendrite pair) was in a consistent direction (+/-) across multiple cells (Fig. 3D). Using a Wilcoxon signed-rank test (p<0.1), we identified 298 genes that met this criterion. For clarity, we will refer to these 298 genes as isoform-specific dendritic (isoDend) RNAs. Most of the isoDend RNAs were categorized as ALEs (249 ALE, 49 tandem), but neither type was significantly enriched in this group compared to the full set of multi-3’UTR genes. Unlike the deDend and consDend sets, the isoDend RNAs were not significantly enriched for particular GO functional categories. Only four of the isoDend RNAs overlapped with the deDend list (*mt-Rnr2, Rpl31, Rpl21*, and *Map2*), indicating that gene-level and isoform-level localized genes are distinct sets. In contrast, approximately half of each the deDend and isoDend sets overlapped with the consDend set (Fig.3E).

Among the 298 isoDend isoform pairs, dendrites preferred the distal isoform in 64% of cases, which was independent of ALE/tandem status. This preference diverged significantly from expectation: in the full set of 3,638 multi-3’UTR genes, dendrites preferred the distal isoform in only 44% of cases (p=3.7e-13; odds ratio=2.4; Fisher’s exact test). Next, we examined the cell-to-cell variability of isoform preferences, particularly focusing on the differences in DF variability between somas and dendrites. For each gene, the variance of DF across samples was calculated separately for soma and dendrite samples. We found that 61.1% of the isoDend genes had a more variable DF in the soma than in the dendrites. Again, this observation diverged significantly from expectation based on the full set of multi-3’UTR genes, where only 29.4% of the genes had a more variable DF in the soma (p<2.2e-16; odds ratio=3.6; Fisher’s exact test).Thus, dendrites showed more specific and consistent isoform preference among the isoDend genes compared to somas, potentially suggesting that certain isoforms are being selectively concentrated in the dendrites due to the presence of *cis* localization signals in the alternative portion of the 3’UTR. Figure 4 provides three representative examples of genes with these isoform patterns, showing the consistent preference for the distal isoform in the dendrites compared to soma for multiple individual cells, and the lower variability of DF in the dendrites compared to the somas. Finally, we looked to see how many of the dendrite-preferred isoforms were among the ~2,000 new, distal 3’UTRs annotated recently by Miura *et al*. in several tissues (Miura et al. 2013). Thirty eight of the dendrite-preferred isoforms overlapped this list (including *Uck2* and *Ube2i* shown in Fig. 4), 12 of which were specific to hippocampal neurons in that study (Miura et al. 2013).

**Figure 4.**
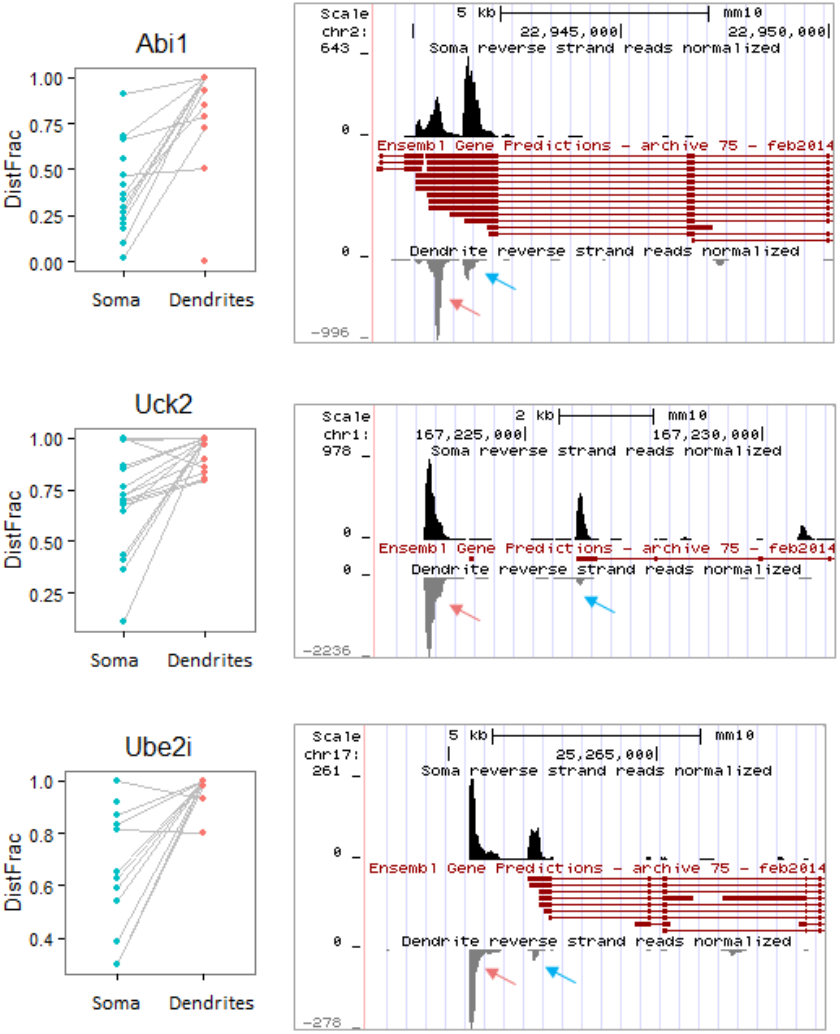
Examples of genes with significantly differentially localized 3’ isoforms. Paired plots on the left show the DF for each soma-dendrite pair (connected by gray lines). The genome browser plots on the right show the read pile-ups for somas (top track; black peaks) compared to dendrites (bottom track; gray peaks; reversed orientation) relative to the annotated gene models from Ensembl (middle track; red). The dendrite-preferred 3’ isoform is indicated by a pink arrow, and the non-preferred isoform is indicated by a blue arrow. Note that for Uck2 and Ube2i, the dendrite-preferred 3’ isoform is a new isoform from (Miura et al. 2013) and thus is not part of the Ensembl gene models. All genes shown are on the reverse strand and thus only reverse-strand reads are displayed.

### Dendritic targeting motifs

We computationally analyzed the deDend, isoDend, and consDend gene lists to identify potential dendritic targeting elements (DTEs) enriched in each set. We first searched for instances of known RBP motifs. The greatest enrichment was seen for SRSF3 binding motif AUCAWCG, which was 2.4 times more common in the deDend RNAs than background and occurred in 59 of the 387 genes in this set. The same SRSF3 motif was also the most enriched motif in the consDend set (1.5 times more common than background) and occurred in 265 of the 1,863 genes in this set. SRSF3 is a brain-expressed splicing factor, and although no specific role for this RBP in neurons has been described, it was recently shown in mouse P19 cells to promote 3’UTR lengthening through distal polyadenylation site usage and promote nuclear export through recruitment of NXF1 (Müller-McNicoll et al. 2016). Therefore, one hypothesis could be that SRSF3 plays a role in the early steps of dendritic localization by promoting inclusion of alternative 3’UTRs (theoretically containing DTEs) and by facilitating nuclear export. We also performed a *de novo* motif analysis using HOMER (Brenner 2010) to see if any previously unidentified motifs were enriched in our sequences. The top motif in each set was UUCGAU (p= 0.0001, odds ratio = 2.9, Hypergeometric test) CCGCAA (p = 1e-7, odds ratio 1.7) and GUGGGU (p = 0.01, odds ratio = 1.2) in the deDend, consDend, and isoDend sets, respectively. One motif, CGCR, was enriched in all three sets, but was only slightly more common in localizers than background (odds ratio < 1.2).

Since G-quaduplexes have been implicated previously in dendritic localization (Subramanian et al. 2011), we also searched our localized sequences for regions that could potentially form this structure. Using a regular expression (see Methods), we searched for potential G-quadruplexes in the 3’UTRs of each localized gene or isoform. G-quaduplexes were 2.0 times more common in the deDend RNAs (p = 0.003, Fisher’s exact test), 1.9 times more common in the consDend RNAs (p = 5.0e-12, Fisher’s exact test), and 1.7 times more common in the isoDend RNAs (not significant; p = 0.14, Fisher’s exact test) than the non-localized background. Overall, 448 of the 2,225 localized genes had at least one potential G-quadruplex in the localized 3’UTR. These results support a possible role for G-quadruplexes in localization in deDend and consDend RNAs, and possibly to a lesser extent in isoDend, but overall it does not appear that this motif alone is enough to explain the majority of localization.

To examine potential structural localization motifs more widely, we applied the *de novo* secondary structure motif-finding tool NoFold (Middleton and Kim 2014) to the localized 3’UTR sequences. Eighty five motifs were significantly enriched compared to non-localized background sequences (p < 0.01, Fisher’s exact test). Two motifs in particular stood out as occurring in a large number of sequences (over 20 unique genes each). Though more conserved on the structure level, the instances of these motifs had enough sequence similarity to suggest a common origin. Using RepeatMasker (Smit, Hubley, and Green 2013), we identified these motifs as instances of the B1 and B2 SINE families, which are ~175nt retrotransposons that form long hairpin structures. To verify that these SINEs were enriched in the localized sequences, we created covariance models (CMs) for B1 and B2 using their canonical sequences and secondary structures and used these CMs to comprehensively identify structurally conserved matches to these elements in our sequences. Compared to non-localized background sequences, B1 structures were found 2.5 times more often in deDend RNAs (p = 0.00047, Fisher’s exact test), 1.8 times more often in consDend RNAs (p = 7.6e-7, Fisher’s exact test), and 1.9 times more often in isoDend RNAs (not significant; p = 0.33, Fisher’s exact test), and B2 structures were found 2.5, 1.9, and 5.7 times more often in the deDend, consDend, and isoDend RNAs respectively (p < 0.001, Fisher’s exact test). Overall, 255 and 165 localized genes out of the 2,225 contained a B1 or B2 match, respectively. These results show that B1 and B2 SINE-related sequences are widespread and over-represented in localized RNAs, suggesting a possible role as DTEs analogous to the role of ID retrotransposon elements in rat dendritic localization (Buckley et al. 2011). Of note, only three genes contained both a G-quadruplex and a B1 or B2 motif, indicating that these signals operate on distinct sets of genes.

### Functional analysis of the “local proteome” using structure information

Only some of the dendritic RNAs might be involved in local protein translation. Nevertheless, to gain a better understanding of potential “local proteome”, we performed a domain-level tertiary structure prediction on the protein products of 1,930 localized mRNAs (combining the deDend, isoDend, and consDend lists and excluding non-coding RNAs). Full length proteins were split into one or more predicted domains (where “domain” is defined as an amino acid chain that likely folds into a compact, independently stable tertiary structure; see Methods), yielding a total of 6,845 domains. Each domain was classified into a SCOP structural fold using our PESS pipeline (Middleton, Illuminati, and Kim 2017). Using this approach, we were able to predict the fold of 2,005 additional domains beyond previous structural annotation (Lees et al. 2012). Using the whole-neuron proteome as a background, we found that the local dendritic proteome was highly enriched for multiple different folds, including several related to cytoskeletal structure such as Spectrin repeats and actin-binding Profilin domains (Fig. 5A). Overall, 503 different folds were represented by at least one domain in the local dendritic proteome, covering almost the entire spectrum of folds expressed in the neuron as a whole (609 folds) (Fig. 5B). This suggests that rather than being highly specialized, the local dendritic RNA has the potential to encode for a diversity of protein functions on par with the whole cell.

**Figure 5.**
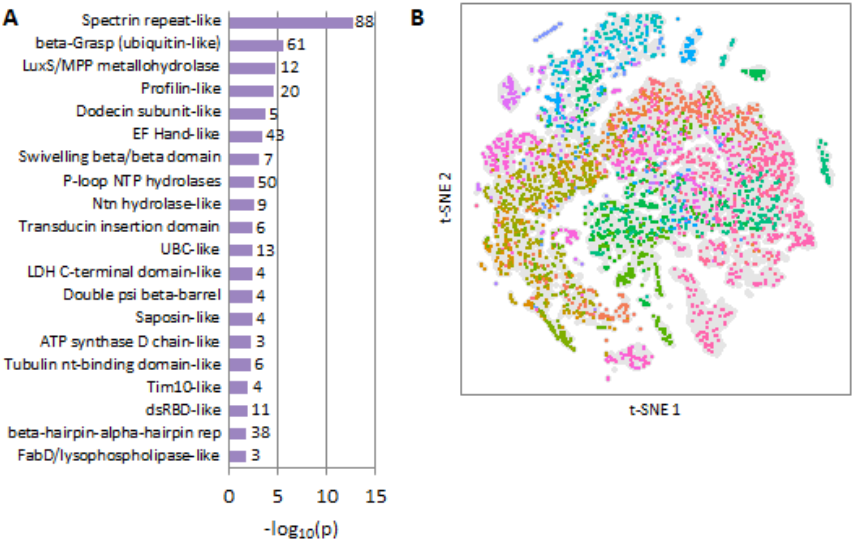
Protein structures of the presumptive locally-translated proteome. (A) SCOP folds enriched in the locally translated proteins compared to the neuron-expressed proteins as a whole. The number of predicted domains in the local proteome for each fold is shown to the right of the bar. (B) Two-dimensional representation of the protein structure space occupied by neuronally-expressed protein domains. All neuronally-expressed protein domains are shown in gray in the background, and locally-translated protein domains are shown in the forefront colored by predicted fold (note that multiple folds may have similar colors due to the large number of folds). Locally translated proteins cover most of the structure space spanned by the whole-neuron set. Projection generated by t-Distributed Stochastic Neighbor Embedding (tSNE) of the PESS coordinates of each input domain.

To highlight some of the insight that can be gained through structure analysis, we selected several folds with important neuronal functions and assessed their representation within the locally translated set, which is described in Supplemental Analysis 1 and Supplemental Tables S2-S4. A full catalog of predicted protein folds is provided in Supplemental Table S5.

### A master list of dendritic RNA

Towards creating a definitive list of dendritic RNAs that have been observed thus far in high-throughput studies, we obtained lists of dendritic genes from six publications that profiled the dendritic transcriptome using microarray or RNA-seq (Ainsley et al. 2014; Cajigas et al. 2012; Lein et al. 2007; Poon et al. 2006; Taliaferro et al. 2016; Zhong, Zhang, and Bloch 2006) and combined those lists with our own. Of a total of 5,635 unique genes on this list, only 1,404 (25%) were observed in at least two studies, and none were found in all studies. The top 40 most frequently observed dendritic genes are listed in Table 1. Ribosomal proteins dominate the list, underscoring the importance of translation-related machinery in the dendrites. The most frequently observed gene was *Tpt1*, a calcium-binding protein involved in microtubule stabilization, which was observed in all but one study. The full list of dendritic genes is available in Supplemental Table S6.

**Table 1.** Top 40 most frequently observed dendritic RNAs.

|  | Gene | # Obs | Refs |
| --- | --- | --- | --- |
| 1 | Tpt1 | 6 | 1,2,4,5,6,7 |
| 2 | Rpl37 | 5 | 1,2,5,6,7 |
| 3 | Rpl4 | 5 | 1,2,4,6,7 |
| 4 | Rps29 | 5 | 1,2,5,6,7 |
| 5 | Rplp0 | 5 | 1,4,5,6,7 |
| 6 | Rpl21 | 5 | 1,2,5,6,7 |
| 7 | Arpc1b | 5 | 1,4,5,6,7 |
| 8 | Ftl1 | 5 | 1,2,4,5,7 |
| 9 | Rps12 | 5 | 1,2,5,6,7 |
| 10 | Ppp1r9b | 5 | 1,2,3,6,7 |
| 11 | Uba52 | 5 | 1,2,5,6,7 |
| 12 | Rpl32 | 4 | 1,5,6,7 |
| 13 | Rpl31 | 4 | 1,2,5,7 |
| 14 | Rpl15 | 4 | 1,2,5,7 |
| 15 | Rpl17 | 4 | 1,2,5,7 |
| 16 | Rpl13 | 4 | 1,2,5,7 |
| 17 | Rpl19 | 4 | 1,2,5,7 |
| 18 | Ids | 4 | 1,2,3,7 |
| 19 | Serbp1 | 4 | 1,2,5,7 |
| 20 | Dlg4 | 4 | 1,2,3,7 |
| 21 | Hint1 | 4 | 1,2,5,7 |
| 22 | Eef2 | 4 | 1,2,4,7 |
| 23 | Rpsa | 4 | 1,2,5,7 |
| 24 | Rps7 | 4 | 1,2,5,7 |
| 25 | Rps2 | 4 | 1,2,5,7 |
| 26 | Rps8 | 4 | 2,5,6,7 |
| 27 | Selenow | 4 | 2,3,5,7 |
| 28 | Rpl36a | 4 | 1,2,5,7 |
| 29 | Pabpc1 | 4 | 1,2,4,7 |
| 30 | Rps25 | 4 | 1,2,5,7 |
| 31 | Rps23 | 4 | 2,5,6,7 |
| 32 | Rps20 | 4 | 1,2,5,7 |
| 33 | Psmc3 | 4 | 1,2,5,7 |
| 34 | Arl3 | 4 | 2,5,6,7 |
| 35 | Eef1b2 | 4 | 1,2,5,7 |
| 36 | Map2 | 4 | 1,2,3,7 |
| 37 | Rplp1 | 4 | 1,2,5,7 |
| 38 | Actb | 4 | 1,2,4,7 |
| 39 | Psd | 4 | 1,2,3,7 |
| 40 | Rpl29 | 4 | 2,5,6,7 |

<sup>1</sup> (Ainsley et al. 2014)
<sup>2</sup> (Cajigas et al. 2012)
<sup>3</sup> (Lein et al. 2007)
<sup>4</sup> (Poon et al. 2006)
<sup>5</sup> (Taliaferro et al. 2016)
<sup>6</sup> (Zhong, Zhang, and Bloch 2006)
<sup>7</sup> This study

## Discussion

Neurons have special RNA localization needs compared to other cell types: their unique morphology—long, extended processes that can be many times the length of the soma— combined with an extensive need for local translation means that neurons must transport a wide variety of RNAs long distances from their origination point in the nucleus. Here, we carried out single neuron sub-cellular RNA sequencing to more precisely identify a total of 2,225 unique genes present in mouse dendrites, including 298 genes for which only a subset of the expressed transcripts were localized, depending on their 3’UTR isoform. Several of these differentially localized 3’UTR isoforms were among the set of recently identified distal 3’UTRs expressed in neurons (Miura et al. 2013). Using *de novo* RNA structure motif analysis, we identified several secondary structures enriched in the 3’UTRs of the localized RNAs, including two hairpin structures derived from B1 and B2 SINE elements, which may act as localization signals. Finally, we applied a protein fold prediction algorithm to make structural and functional predictions for the set of proteins that are putatively translated locally at the synapse.

Based on our results, there are almost 300 genes with alternative 3’ isoforms where one isoform was consistently more dendritically localized than the other. The use of alternative 3’UTRs is an attractive model for how neurons might regulate localization, especially since 3’UTRs theoretically have the potential to provide an element of tissue-specificity to localization. In light of this, it is somewhat surprising that of the 38 dendrite-targeted isoforms we identified that were also profiled by (Miura et al. 2013), only 12 were specific to hippocampal neurons according to the Miura data. The other 26 isoforms were found in at least one of the other mouse tissue types profiling in that study, which included spleen, liver, thymus, lung, and heart, suggesting a general lack of tissue-specificity of these dendritically-targeted isoforms. Instead, we postulate that tissue-specific localization may be achieved by tissue-restricted expression of *trans* factors (e.g. RBPs) rather than by regulation of DTE-containing isoform expression. In addition, although we observed significant enrichment of several candidate DTEs, including RBP recognition sites, G-quaduplexes, and SINE mediated hairpin structures, none of the potential regulatory elements were universal nor unique to localized RNA sequences. These results suggest that dendritic RNA localization involves multiple pathways and overlapping mechanisms (Buckley et al. 2011; Holt and Schuman 2013), and that “aggregate” localization signals composed of multiple DTEs may be necessary to improve specificity and possibly also refine the destination of dendritically targeted transcripts.

An intriguing finding was that the composition of the deDend set was skewed towards RNAs that encode proteins that modulate RNA translation and mitochondrial function, as compared to the larger consDend set which covered many more dendrite-and synapse-specific functions. This leads us to speculate that translational regulation of dendritic protein synthesis might be dynamically modulated through stimulated transient local production of proteins that enhance the capacity to make ATP and to “jump start” the translational machinery. This jump start model would postulate a generalized but specific regulatory mechanism that could act on whatever RNAs are present at the site, without the need for individualized translation regulation of each dendritic RNA. Such a mechanism would allow the standard cellular translation mechanism to be specific without requiring the existence of new RNA transport proteins or transcript-specific translation. Regulation of local protein synthesis by the global mechanism of spatial translational control as opposed to individual RNA translational enhancement is different from current models of how dendritic protein synthesis is regulated, suggesting avenues for future experiments.

A crucial remaining question is what role individual locally translated proteins play in long-lasting synaptic potentiation. The post-synaptic density and surrounding dendritic spine are highly structured formations that depend on a scaffold of interacting proteins (Kim and Sheng 2004; Dalva, McClelland, and Kayser 2007; Zheng et al. 2011), which in turn usually require a specific three-dimensional fold in order to function properly. Here, we provide a fold-level structure-function annotation of 1,930 proteins that we predict to be locally translated at the synapse based on our RNA localization analysis. Given that mutations linked to neuropsychiatric diseases have been found to be enriched in synaptic proteins in human and mouse, and several of these mutations appear to disrupt important structures (Liu-Yesucevitz et al. 2011; Grant 2012), structural knowledge of these proteins is important for understanding these disorders. A more complete picture of the structures of locally translated proteins will help both in functional understanding and mutation-impact analysis.

One limitation of our study is that neurons were only surveyed at the basal state, rather than after synaptic stimulation. Several studies have shown that RNA localization changes after stimulation (Tongiorgi, Righi, and Cattaneo 1997; Steward et al. 1998; Eberwine et al. 2001; Yoon et al. 2016); therefore, the set of dendrite RNAs identified here may still be only a subset of the RNAs needed for LTP. There also may be important differences between neurons in culture and *in vivo* that would be missed in our analysis. We observed significant overlap between our localized set and a set of localized RNAs derived partly from tissue-based studies conducted after fear conditioning (Ainsley et al. 2014), suggesting a reasonable amount of concordance between basal primary cultures and post-stimulation tissue samples. Nonetheless, an important future direction will be to repeat the sub-cellular sequencing described here after stimulation. It will be particularly interesting to see if groups of RNAs that share a DTE undergo coordinated changes in localization post-activation, and conversely, if coordinated RNAs share any new DTEs.

In sum, our study represents a comprehensive resource for RNA localization in mouse neurons consisting of our new sub-cellular RNA sequencing dataset, a compilation of previous dendritic RNA studies, as well as computational annotation of motifs and structures. The resource generated here may have broad utility for continued study of mechanisms of dendritic RNA localization and the role of localized RNA in neuronal function and dysfunction.

## Materials and Methods

### Neuron culture and collection

Hippocampal neurons from embryonic day 18 (E18) mice (C57BL/6) were cultured as described in (Buchhalter and Dichter 1991) for 15 days. Isolated single neurons were selected for collection. A micropipette with a closed, tapered end was used to sever dendrites from the cell body. Another micropipette was used to aspirate the soma, which was deposited into a tube containing first strand synthesis buffer and RNase inhibitor and placed on ice. A separate micropipette was used to aspirate the dendrites, which were deposited into a separate tube as above. Samples were transferred to −80°C within 30 minutes and stored there until first strand synthesis. Sixteen neurons (32 total samples) were collected from multiple cultures across multiple days.

### Single cell RNA amplification and sequencing

ERCC spike-in control RNA was diluted 1:4,000,000 and 0.9uL was added to each tube. Poly-adenylated RNA was amplified using two or three rounds of the aRNA *in vitro* transcription-based amplification method, as described in (Morris, Singh, and Eberwine 2011). The quality and quantity of the amplified RNA was verified using a Bioanalyzer RNA assay. Strand-specific sequencing libraries were prepared using the Illumina TruSeq Stranded kit according to the manufacturer’s instructions, except that the initial poly-A capture step was skipped because the aRNA amplification procedure already selects for poly-adenylated RNA. Samples were sequenced on a HiSeq (100bp paired-end) or NextSeq (75bp paired-end) to an average depth of 25 million reads. Reads were trimmed for adapter and poly-A sequence using in-house software and then mapped to the mouse genome (mm10) using STAR (Dobin et al. 2013). Uniquely mapped reads were used for feature quantification using VERSE (Zhu et al. 2016). The features used for each analysis are described below.

### Gene-level expression and localization

Three sources of gene annotations were combined to obtain a comprehensive definition of known 3’ ends: Ensembl genes (downloaded from UCSC, Dec 2015); UCSC genes (downloaded from UCSC, Dec 2015); and the set of ~2,000 new 3’UTRs determined by Miura et al. (Miura et al. 2013). The 3’UTR regions of these annotations were used for quantification of reads. A single 3’UTR feature was created for each gene by taking the union of all 3’UTR regions for that gene. Read counts were calculated for each gene based on how many reads mapped to this 3’UTR region. Quantification was done using VERSE with options “-s 1 -z 3 -- nonemptyModified”. For differential expression analysis, we used only the genes that had at least one read in at least half (16) of the samples. Read counts were normalized and differentially expressed genes between the dendrites and soma were identified using DESeq2 with a paired experimental design. A FDR corrected p ≤ 0.05 was used to identify significantly differentially expressed genes. The consDend genes were identified separately based on having at least 1 read in at least 90% (i.e. 15 out of 16) of the dendrite samples.

GO functional enrichment of deDend and consDend genes was calculated using the GOrilla webserver (Eden et al. 2009). For deDend genes, the background set for GO analysis was all genes with at least one read in half the samples; for the consDend genes, the background was all genes with at least one read in at least 15 samples (i.e. the input sets for each analysis).

Gene markers of pyramidal neurons and cardiomyocytes, as well as housekeeping genes, were obtained from (Dueck et al. 2015). Markers of other mouse brain cell types were obtained from (Zhang et al. 2014).

### Isoform-level expression and localization

To quantify individual 3’ isoforms of genes, we used the last 500nt of each 3’ end for that gene as the isoform quantification feature. Any 3’ ends that were less than 500nt apart were merged together into a single quantification feature. Thus, the final set of 3’ isoform quantification features is non-overlapping. Isoform read counts were calculated by VERSE using the same parameters as above. Genes with only one expressed 3’ isoform were removed from further analysis to focus on alternative expression of 3’ isoforms.

To identify the top two 3’ isoforms for each gene, the following procedure was used. For each gene in each sample, the fraction of reads mapping to each isoform was calculated (that is, the number of reads mapping to that isoform divided by the total reads for all isoforms of the gene). The fractions for each isoform were then summed up across samples (unless a sample had fewer than 10 reads total for that gene, in which case it was skipped) and the two isoform with the highest total per gene were considered the top two isoforms for that gene. The purpose of this process was to give each sample equal weight in the final decision of the top 3’UTR, while also excluding samples with too few reads to give a reliable estimate of the isoform fractions. This process was repeated for each gene with at least two expressed isoforms in the dataset. Then for each gene, whichever of the top two isoforms was more 5’ (as defined by the locations of their 500nt quantification features) was designated the “proximal” isoform, and whichever was more 3’ was designated the “distal” isoform. Finally, for each gene in each sample, we calculated the distal fraction (DF) as the fraction of reads mapping to the distal isoform divided by the total reads mapping to the distal and proximal isoforms.

We defined the proximal and distal isoforms as being, relative to each other, generated by alternative splicing (ALEs) or alternative cleavage and polyadenylation (Tandem UTRs) by the following criterion: if the full length 3’UTRs of a pair of isoforms were directly adjacent or overlapping, they were called tandem; otherwise, they were called ALEs.

The differential localization of isoforms was determined based on the change in distal fraction between soma and dendrites of the same original neuron. A non-parametric paired test of differences (Wilcoxon signed-rank test) was used to identify genes with consistent changes in distal fraction across samples. Only genes with at least five pairs of samples (where a “pair” means the soma and dendrites from the same original neuron) where each member of the pair had at least 10 combined reads for the two isoforms were tested (3,638 genes), to ensure there was enough read-and sample-support to reliably identify these events.

GO enrichment was done on the dendrite-enriched isoforms as described in the previous section, using the input set of 3,638 genes as background.

### Background datasets for motif enrichment

We generated a pool of “non-localized” background sequences based on the list of genes that were significantly higher expressed in the soma from the gene-level DESeq2 analysis described above. We filtered this set to remove any overlap with one of the other localized lists (i.e. the consDend list and the isoDend list) and any overlap with previously annotated dendritically localized genes in order to make this list as specific to non-localized genes as possible. Since motif frequency in a sequence can be related to sequence length, we created a length-matched background set for each of the three localized gene lists as follows: (1) for each localized gene in the set, scan the pool of non-localized genes in order of their somatic specificity (starting with the most soma-specific, as indicated by its DESeq2 test statistic); (2) select the first non-localized gene encountered with a 3’UTR length within 100nt of the localized gene’s 3’UTR length; (3) add the selected non-localized gene to the background set and remove it from the pool; (4) if no background gene can be found that meets the 100nt criteria, select whichever gene in the pool that has the most similar 3’UTR length to the localized gene’s 3’UTR. Using this protocol resulted in background sets with highly similar length characteristics to the foreground set.

### RNA motif analysis

Linear motifs were identified using the HOMER motif-finding suite (Brenner 2010). *De novo* enriched motif searches were done using the script “findMotifs.pl” and set to look for either short motifs (4 or 6nt) or long motifs (8, 10, or 12nt). Enrichment of known RBP binding motifs was analyzed using the same script with option “-known” in combination with a custom set of positional weight matrices specifying binding preferences that was downloaded from CISBP-RNA (version 0.6) (Ray et al. 2013). A log-odds threshold for RBP motif matching was set for each motif separately based on the number of informative positions in the motif such that longer, more specific motifs had a higher log-odds threshold for calling a match. The background sets used for enrichment testing were the length-matched non-localized sets described above.

G-quadruplexes were identified by regular expression search using the “re” module in Python. The search pattern was ‘([gG]{3,}\w{1,7}){3,}[gG]{3,}’, which requires three consecutive matches to the pattern “three or more G’s followed by 1-7 of any nucleotide” and then ending with a fourth set of three or more G’s. The background set was the same as described in the previous section.

*De novo* identification of enriched RNA secondary structures was performed using NoFold (Middleton and Kim 2014). Sliding windows of 100nt (slide = 75nt) across the localized sequences were used for input. Background datasets were the same as described in the previous section and also converted to sliding windows with the same parameters. Additional matches to the B1 and B2 elements were found by creating a CM for each element based on its canonical sequence(s) downloaded from RepeatMasker (Smit, Hubley, and Green 2013) and its predicted MFE structure from RNAfold (Gruber et al. 2008). The sequences and structures used to create the CM are as follows:

B1 sequence:

GAGGCAGGCGGATTTCTGAGTTCGAGGCCAGCCTGGTCTACAGAGTGAGTTCCAGGACAGCCAGGGCTACACAGAGAAACCCTGTCTC

B1 structure:

((((((((….(((((((((((..(((…(((((.((……..))..)))))…))).)))))…))))))…))))))))

B2 sequence:

GCTGGTGAGATGGCTCAGTGGGTAAGAGCACCCGACTGCTCTTCCGAAGGTCAGGAGTTCAAATCCCAGC

B2 structure:

(((((.((..((((((….((.(((((((……)))))))))………))).)))..)))))))

Bitscore cutoffs for high-quality matches were set to 50 for B1 and 35 for B2 based on the length of the model. Enrichment was computed using Fisher’s exact test based on the number of high quality matches in the localized set compared to the non-localized background (same background as above). Only one match was counted per gene for the purposes of enrichment testing.

### Protein structure analysis

For each predicted dendritic RNA we obtained the canonical protein sequence, if any, from UniProt (The UniProt Consortium 2017). The canonical isoform is defined by UniProt to usually be the one that is most inclusive of exons/domains. We refer to this protein set as the “local proteome”. We also obtained the canonical protein sequences for the full set of expressed genes in soma and dendrite samples (at least 1 read in at least 15 samples) to use as a background for comparison with the local proteome.

Each protein was split into domains based on DomainFinder Gene3D predictions (Yeats, Redfern, and Orengo 2010; Lees et al. 2012). If there were regions between, before, or after predicted domains that were longer than 30 amino acids (aa) but did not have a Gene3D prediction, we also included these. If a “filled in” region such as this was longer than 450 aa, we used a sliding window of 300 aa (slide = 150 aa) to break it into smaller pieces, since domains are rarely larger than this. The fold of each domain was predicted using the method described in (Middleton, Illuminati, and Kim 2017). A nearest neighbor distance threshold of ≤ 17.5 was used to designate “high confidence” predictions, and a more lenient threshold of ≤ 30 was used to designate “medium confidence” predictions.

## Acknowledgements

This work was funded in part by NIMH U01MH098953 to JK and JE, NIGMS R01 GM110005 to JE and JK, and Health Research Formula Fund from the Pennsylvania Commonwealth to JK. SM was supported by a DOE CSGF fellowship (DE-FG02-97ER25308). The funding agencies played no direct role in design, analyses, and conclusions presented in this work.

